# Acyl-coA binding protein AcbdA regulates peroxisome hitchhiking on early endosomes

**DOI:** 10.1101/2025.04.17.649231

**Authors:** Bellana Driscoll, Madison B. Fountain, Isabella N. Gates, Reihane Abdollahi, Allison M. Langley, Matthew B. Owens, Jenna R. Christensen, John Salogiannis

**Affiliations:** Department of Molecular Physiology and Biophysics, University of Vermont Larner College of Medicine, Burlington, VT 05405; Department of Molecular Biosciences, Northwestern University, Evanston IL 60208; Neuroscience Graduate Program, University of Vermont, Burlington, VT 05405; Cellular, Molecular and Biomedical Sciences Program, University of Vermont, Burlington, VT 05405

## Abstract

Motor-driven transport on microtubules is critical for distributing organelles throughout the cell. Most commonly, organelle movement is mediated by cargo adaptors, proteins on the surface of an organelle that directly recruit microtubule-based motors. An alternative mechanism called hitchhiking was recently discovered: some organelles move, not by recruiting the motors directly, but instead by using membrane contact sites to attach to motor-driven vesicles and hitchhike along microtubules. Organelle hitchhiking is observed across fungi and animals. In filamentous fungi, nearly all peroxisomes move by hitchhiking on early endosomes (EEs). In the fungus *Aspergillus nidulans*, EE-associated linker proteins PxdA and DipA are critical for establishing EE-peroxisome membrane contact sites required for peroxisome movement. How peroxisomes recognize this subset of EEs and what peroxisome-membrane proteins exist that can interact with EEs is not known. Here, we undertook a forward mutagenesis screen to identify such proteins. We discovered an acyl-coA binding (ACB) domain-containing protein AcbdA/AN1062 that localizes to peroxisomes via its tail-anchored transmembrane domain (TMD). Deleting the AcbdA gene or only its N-terminal ACB domain perturbs the movement and distribution of peroxisomes. Importantly, AcbdA is not required for the movement of EEs or for the recruitment of PxdA and DipA on EEs. Fatty acid (FA)-induced increases in peroxisome movement require AcbdA, suggesting that peroxisome hitchhiking on EEs is coupled to FA metabolism. Mutating a conserved FFAT motif, predicted to interact with the endoplasmic reticulum (ER), has no effect on peroxisome movement. Taken together, our data indicate that AcbdA is a peroxisome-membrane protein required to tether peroxisomes to EEs during hitchhiking. AcbdA’s involvement in peroxisome-EE contact site formation represents a divergence from known functions of Acbd4/5 proteins and adds layers to our understanding of the functionality of the Acbd4/5 family of proteins.

## Introduction

Microtubule-based transport of cargos (organelles, vesicles, and macromolecular complexes) is critical for a wide-variety of cellular functions (Barlan and Gelfand, 2017). Cargos are driven by molecular motors dynein and kinesins to ensure proper localization at the right time and place. Canonically, most cargos move by directly recruiting motors via cargo adaptors (Cianfrocco *et al*., 2015; Hoogenraad and Akhmanova, 2016; Reck-Peterson *et al*., 2018; Cross and Dodding, 2019; Olenick and Holzbaur, 2019; Xiang and Qiu, 2020). These adaptors usually interact with only a select few cargos, a mechanism that helps ensure cargo specificity during transport.

For example, Hook family protein adaptors associate with endosomes to recruit dynein and/or kinesin (Bielska *et al*., 2014; Zhang *et al*., 2014; Kendrick *et al*., 2019; Siddiqui *et al*., 2019; Carvalho *et al*., 2025), Trak1/2 proteins associate with mitochondria (Glater *et al*., 2006; Wang and Schwarz, 2009; van Spronsen *et al*., 2013; Fenton *et al*., 2021), and BicD2 associates with Golgi-derived vesicles marked by the GTPase Rab6 (Splinter *et al*., 2012; Hoogenraad and Akhmanova, 2016; Huynh and Vale, 2017).

A second prominent mode of transport called ‘hitchhiking’ has recently emerged. During hitchhiking, some cargos move not by recruiting adaptors and motors directly, but instead by tethering and co-transporting with motor-driven vesicles (Salogiannis and Reck-Peterson, 2017; Christensen and Reck-Peterson, 2022). In this way, cargo movement is coupled to moving vesicles rather than to molecular motors. In the pathogenic filamentous fungus *Ustilago maydis*, macromolecular complexes such as septin complexes, mRNA-ribonucleoprotein complexes, and ribosomes hitchhike on the surface of Rab5-marked early endosomes (EEs) (Baumann *et al*., 2012, 2014; Higuchi *et al*., 2014; Haag *et al*., 2015; Pohlmann *et al*., 2015; Zander *et al*., 2016). mRNA hitchhiking is also found in other systems including budding yeast and mammalian neurons (Aronov *et al*., 2007; Genz *et al*., 2013; Liao *et al*., 2019; De Pace *et al*., 2024).

More recently, it was observed that hitchhiking occurs at membrane contact sites (MCS) between membrane-bound organelles and Rab-marked vesicles. In the filamentous fungus *Aspergillus nidulans*, peroxisomes tether to and are pulled along microtubules on Rab5-marked EEs (Salogiannis *et al*., 2016). In the filamentous fungi *U. maydis*, peroxisomes, lipid droplets, and ER all hitchhike on EEs (Guimaraes *et al*., 2015). In both instances, EEs move by interacting with motors dynein (NudA) and kinesin-3 (UncA) scaffolded by a cargo adaptor complex HookA/Hok1 (Wedlich-Söldner *et al*., 2002; Abenza *et al*., 2009; Zekert and Fischer, 2009; Schuster *et al*., 2011; Bielska *et al*., 2014; Yao *et al*., 2014; Zhang *et al*., 2014). Perturbing the motility of EEs abolishes movement of hitchhiking organelles and leads to their aberrant accumulation at the hyphal tip (Zhang *et al*., 2014; Guimaraes *et al*., 2015; Salogiannis, Christensen *et al*., 2021). In mammalian cells, peripheral tubules of the ER hitchhike on multiple Rab-marked vesicles, including ER-to-Golgi vesicles marked by Rab1, lysosomes and endolysosomal vesicles marked by Rab5 and Rab7, and Golgi-derived vesicles marked by Rab6 (Friedman *et al*., 2013; Guo *et al*., 2018; Lu *et al*., 2020; Spits *et al*., 2021; Jang *et al*., 2022; Langley, Abeling-Wang *et al*., 2025). Taken together, organelle hitchhiking at MCS is a novel form of microtubule-based transport that is more common than previously appreciated. The molecular mechanisms leading to loading and offloading of hitchhiking cargos, the biological function of these contacts, and the tethers involved are not well understood.

Peroxisome hitchhiking on EEs in filamentous fungi is one of the best-studied examples of organelle hitchhiking. Peroxisomes are metabolic organelles with diverse functions including β-oxidization of very long chain fatty acids (VLCFAs), secondary metabolite synthesis, and scavenging of reactive oxidative species (Maruyama and Kitamoto, 2013; Steinberg, 2016). How and why peroxisomes move is not well understood. While the peroxisome-EE tethers in *U. maydis* have not been identified, in *A. nidulans*, peroxisome hitchhiking requires the EE-associated linker proteins PxdA and DipA (Salogiannis *et al*., 2016, Salogiannis, Christensen *et al*., 2021). Depleting either protein renders peroxisomes immotile, but EEs move normally, suggesting that the role hitchhiking linkers play is to regulate peroxisome attachment to EEs. How peroxisomes recognize and contact EEs during movement is not known (Figure 1A).

**Figure 1:**
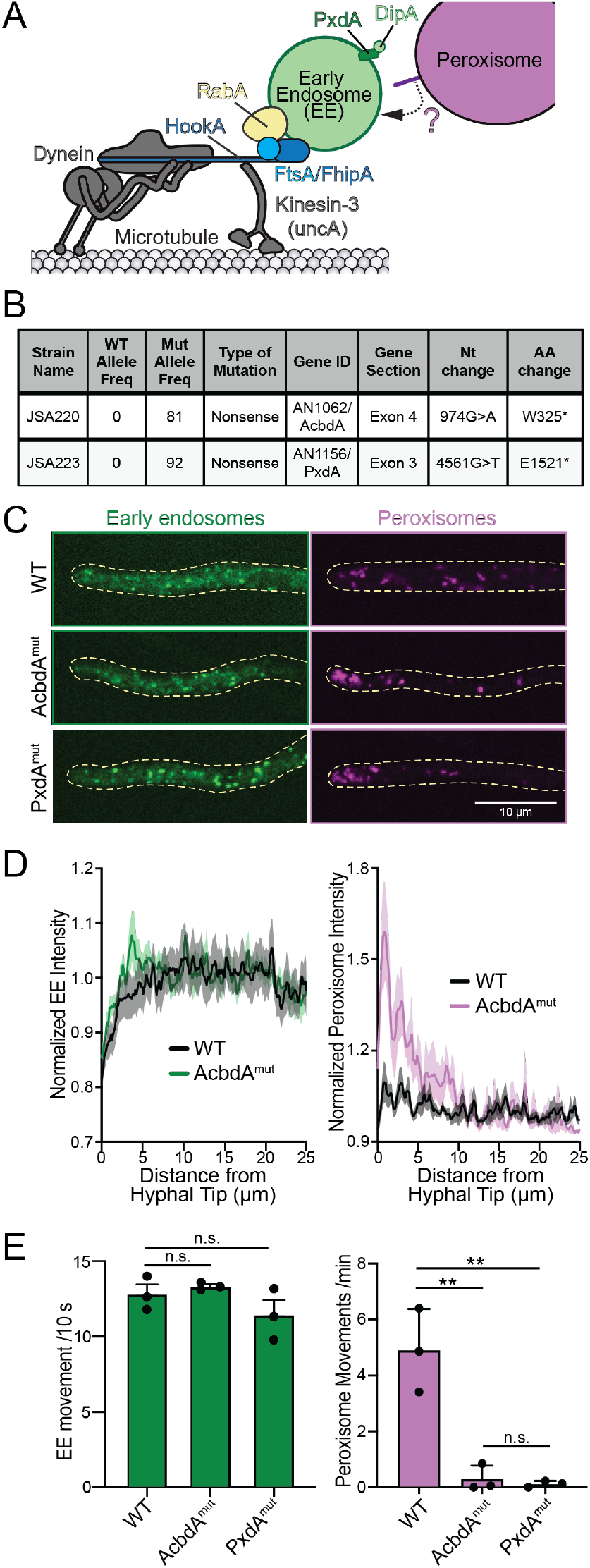
Mutagenesis screen reveals AN1062/AcbdA as a regulator of peroxisome movement. (**A**) Hitchhiking model; peroxisome-localized proteins mediating peroxisome hitchhiking on EEs is unknown. (**B**) Summary table of mutagenized strains whole genome sequencing results. (**C**) Micrographs of EEs (RabA-GFP) and peroxisomes (mCherry-PTS1) along hyphae from wild-type or mutagenized strains. (**D**) Fluorescence intensity line scans on a single optical section from hyphal tip of EEs (left) and peroxisomes (right); (*p<0.05; two-way ANOVA, Bonferroni post hoc multiple comparisons test significant from 0.1083 to 1.5167 µm and 2.2750 to 3.1417 µm; n=12 (WT) and 7 (AcbdA^mut^) hyphae for EEs and 10 (WT) and 8 (AcbdA^mut^) hyphae for peroxisomes). Note: y-axis starts at 0.7 for EEs, and 0.9 for peroxisomes. (**E**) Bar graphs of quantified EE (left) and peroxisome (right) movements (mean ±S.D.) near the hyphal tip of WT and mutagenized strains. Mean EE movements are 12.82 ±1.10 for WT, 13.34 ±0.23 for AcbdA^mut^ and 11.44 ±1.69 for PxdA^mut^ hyphae. (STATS). Mean peroxisome movements are 4.90 ±1.49 for WT, 0.30 ±0.47 for AcbdA^mut^ and 0.12 ±0.11 for PxdA^mut^ (*p<0.01; one-way ANOVA, Dunnett’s post hoc multiple comparisons test; N=3 technical replicates from 44-53 hyphae per condition). Also see Supplemental Movie S1.

Here, we performed a mutagenesis screen in *A. nidulans* to identify novel regulators of peroxisome hitchhiking on EEs, specifically focused on identifying peroxisome-associated membrane tethers that facilitate contact with EEs. We discovered an uncharacterized acyl-coA binding (ACB) domain-containing protein AcbdA. AcbdA is a member of the Acbd4/5 family of proteins and shares some similarities and differences with other characterized Acbd4/5 proteins. AcbdA localizes to peroxisomes via a tail-anchored transmembrane domain and requires its N-terminal ACB domain to regulate the movement of peroxisomes. Importantly, AcbdA does not regulate the movement of EEs or the localization of PxdA and DipA. AcbdA is critical for the link between FA metabolism and peroxisome movement. Additionally, though AcbdA has a FFAT motif similar to other Acbd4/5 proteins, the FFAT motif is not required for peroxisome motility. Overall, we have identified a novel member of the Acbd4/5 family of proteins as critical for peroxisome hitchhiking on EEs.

## Results and Discussion

### A forward mutagenesis screen identifies the acyl-coA binding domain-containing AcbdA/AN1062

We set out to identify regulators of peroxisome hitchhiking in *A. nidulans* using a forward genetic screen. To accomplish this, we subjected a strain with fluorescently-labeled peroxisomes (mCherry-Flag-PTS1) and EEs (mTagGFP-RabA/5a; see Supplemental Figure S1A for table of strains used in this study) to the mutagen 4-Nitroquinoline 1 oxide (4NQO) at a concentration resulting in colony survival rates of 10-30% (1.3µg/mL; Supplemental Figure S1B). This concentration was used previously to identify PxdA, as well as other regulators of microtubule-based transport (Tan *et al*., 2014; Salogiannis *et al*., 2016). We performed a visual-based microscopy screen (∼8,000 colonies) for defects in the movement and distribution of peroxisomes and EEs (see Methods). We hypothesized that strains with defective peroxisome distribution but normal EE distribution would likely have mutations in genes involved in MCS formation between EEs and peroxisomes, as known regulators of peroxisome hitchhiking do not affect EE motility (Salogiannis *et al*., 2016, Salogiannis, Christensen *et al*., 2021; Songster *et al*., 2023). Therefore, colonies exhibiting defective distribution of peroxisomes but normal distribution of EEs were scored as positive hits. We did not screen colonies with marked growth defects as these strains would likely yield mutations in genes affecting nuclear distribution or developmental defects (Xiang *et al*., 1994, 1999). We also did not follow up on strains with reduced numbers of peroxisomes, as those would likely encode peroxins or proteins involved in peroxisome biogenesis (Platta and Erdmann, 2007; Hynes *et al*., 2008).

We identified two mutagenized strains with distribution defects in peroxisomes, but not EEs (Figure 1, B and C). Both strains had immotile peroxisomes that were accumulated at the hyphal tip (Figure 1, B – D; Supplemental Movie S1). In contrast, the movement and distribution of GFP-RabA/5a marked EEs were normal (Figure 1, C – E; Supplemental Movie S1). A backcrossing strategy followed by whole genome sequencing was employed to identify high-quality mutations (see Methods; Tan *et al*., 2014). Only one phenotypic-causing mutation was identified for each strain (Figure 1B). Both strains contained nonsense mutations: one mapped to the previously identified hitchhiking linker PxdA/AN1156. The other strain mapped to a gene encoding an acyl-coA binding (ACB) domain-containing protein AN1062 (Figure 1B), which we renamed AcbdA and further characterized in this study.

### AcbdA regulates the movement and distribution of peroxisomes but not early endosomes

We hypothesized that AcbdA represents a new regulator of peroxisome hitchhiking. To begin to address this, we monitored peroxisomes in an *acbdA* deletion strain (*ΔacbdA*; Supplemental Figure S1C). Similar to the mutagenized strain from our screen, peroxisomes were drastically accumulated at the hyphal tip in the *ΔacbdA* strain compared to WT (Figure 2, A and B). Furthermore, the majority of peroxisomes were immotile in *ΔacbdA* hyphae (Figure 2C; Supplemental Movie S2). In those that moved, the speed and distance traveled (i.e., run length) were also reduced (Figure 2, D and E). Overall, the peroxisome defects observed in *ΔacbdA* strains phenocopied *ΔpxdA* strains (Salogiannis *et al*., 2016).

**Figure 2.**
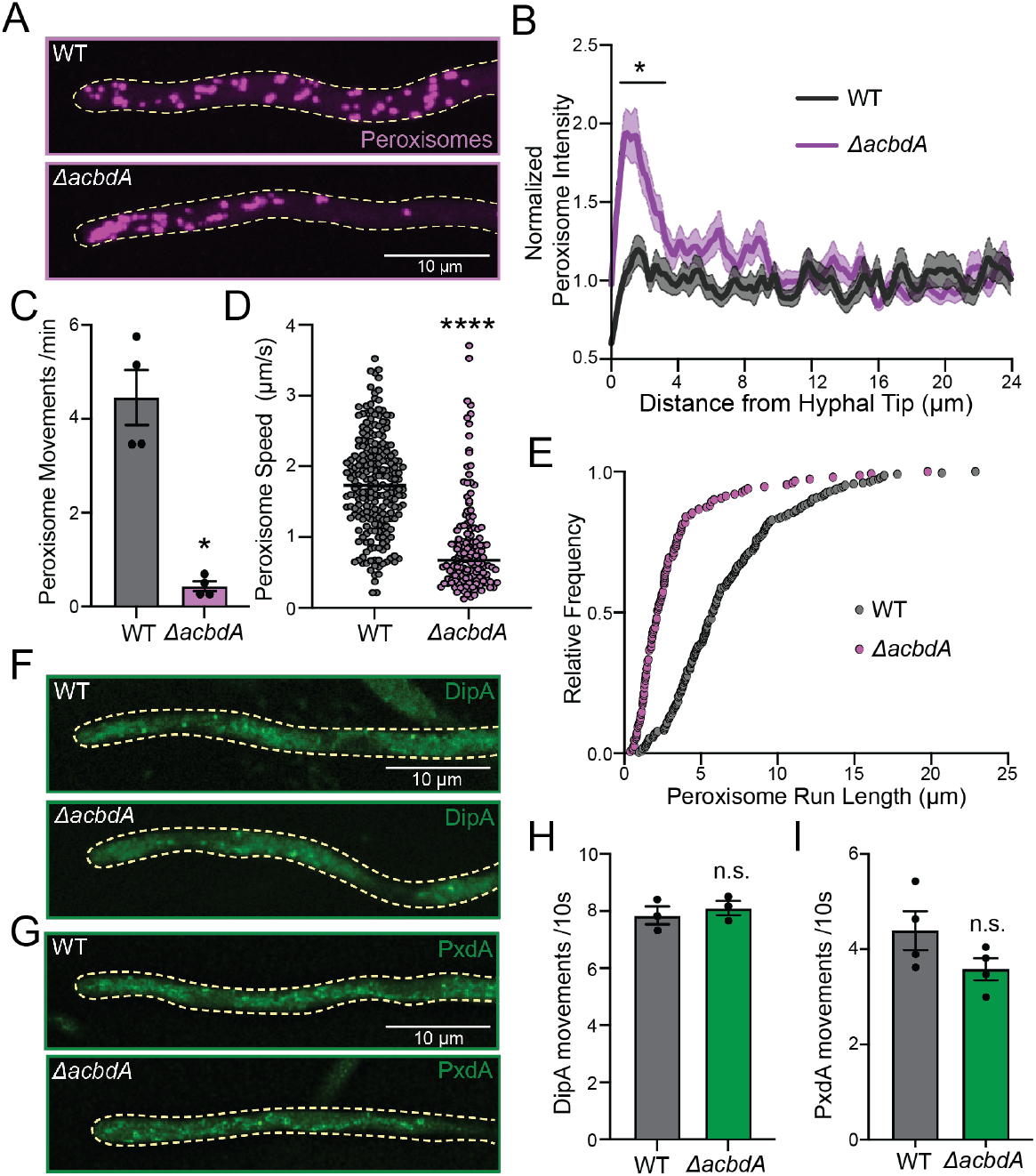
AcbdA regulates the movement of peroxisomes, but not PxdA- or DipA-marked EEs. (**A**) Representative micrographs of WT and *ΔacbdA* hyphae expressing fluorescently labelled peroxisomes (mCherry-PTS1). (**B**) Peroxisome distribution quantified from line scans of fluorescent micrographs from z-stacks of germling hyphae. Fluorescence intensity as a function of distance from the hyphal tip is displayed as mean (solid lines) ±SEM (shading). Distribution near the tip was significantly different between WT and *ΔacbdA* (*p<0.05; two-way ANOVA, Bonferroni’s multiple comparisons test significant between 0.43 µm and 2.38 µm; n=73 (WT) and 78 (*ΔacbdA*) hyphae). Note y-axis starts at 0.5. (**C**) Bar graph of peroxisome movements per min. Peroxisome movements are 4.46 ±0.59 for WT and 0.43 ±0.10 for *ΔacbdA* (*p<0.05; Mann-Whitney test; N=4 technical replicates from 54 (WT) and 45 (*ΔacbdA*) hyphae). Also see Supplemental Movie S2. (**D**) Scatter plot of instantaneous speeds of moving peroxisomes. Mean WT speed of peroxisomes was 1.73 µm/s and *ΔacbdA* speed was 0.67 µm/s (****p<0.0001; Student’s t-test; n=228 (WT) and 150 (*ΔacbdA*) moving peroxisomes). (**E**) Cumulative distribution of run lengths of moving peroxisomes. The decay constant (τ) is 7.96 for WT and 3.11 for *ΔacbdA* (n=228 (WT) and n=150 (*ΔacbdA*) run lengths). (**F** and **G**) Representative micrographs of WT and *ΔacbdA* hyphae expressing fluorescently labelled (F) DipA (DipA-2xmTagGFP) and (G) PxdA (PxdA-mTagGFP). (**H**) Bar graph of DipA movements in WT and *ΔacbdA* hyphae. DipA movements are 7.86 ±0.31 for WT and 8.11±0.24 for *ΔacbdA* (p=0.70 [n.s]; Mann-Whitney test; N=4 technical replicates from 17 (WT) and 18 (*ΔacbdA*) hyphae). Also see Supplemental Movie S2. (**I**) Bar graph of PxdA movements in WT and *ΔacbdA* hyphae. Mean PxdA movements are 4.40 ±0.40 for WT and 3.58 ±0.23 for *Δacbd*A (p=0.20 [n.s.]; Mann-Whitney test; N=4 technical replicates from 69 total hyphae for both genotypes). Also see Supplemental Movie S2. Movements from (C, H, and I) quantified near the hyphal tip. All error bars SEM.

Our screen determined that a mutation in AcbdA does not affect the distribution and movement of Ees (Figure 1, C - E). One possibility is that AcbdA recruits or regulates known EE-localized hitchhiking proteins PxdA and DipA. Depleting these proteins has previously been shown to have no effect on RabA-marked EEs (Salogiannis *et al*., 2016; Salogiannis, Christensen *et al*., 2021). To test this, the movement of PxdA and DipA was quantified in *ΔacbdA* strains. No differences in movement or distribution of PxdA or DipA were observed compared to a WT control (Figure 2, F - I; Supplemental Movie S2). This finding suggests that AcbdA is not involved in recruiting known hitchhiking regulators or in regulating EE movement.

### AcbdA localizes to peroxisomes via its tail-anchored transmembrane domain

AcbdA is a 384 aa protein with an N-terminal ACB domain, an internal two phenylalanines in an acidic tract (FFAT)-motif, an internal coiled-coil, and a C-terminal transmembrane domain (TMD) (Figure 3A). BLASTp searches using AcbdA as the query identified other proteins with similar domain architecture, including known members of the Acbd4/5 family of proteins (Figure 3A). Therefore, we suspect that AcbdA is a member of the Acbd4/5 family of proteins (Islinger *et al*., 2020). Other characterized Acbd4/5 proteins such as those from humans (Hs_ACBD4 and Hs_ACBD5), *Drosophila melanogaster* (Dm_Acbd4/5) and *U. maydis* (Um_Acbd4/5) share features with AcbdA, including an N-terminal ACB domain and a C-terminal TMD (Figure 3A), consistent with a recent study (Kors *et al*., 2024).

**Figure 3.**
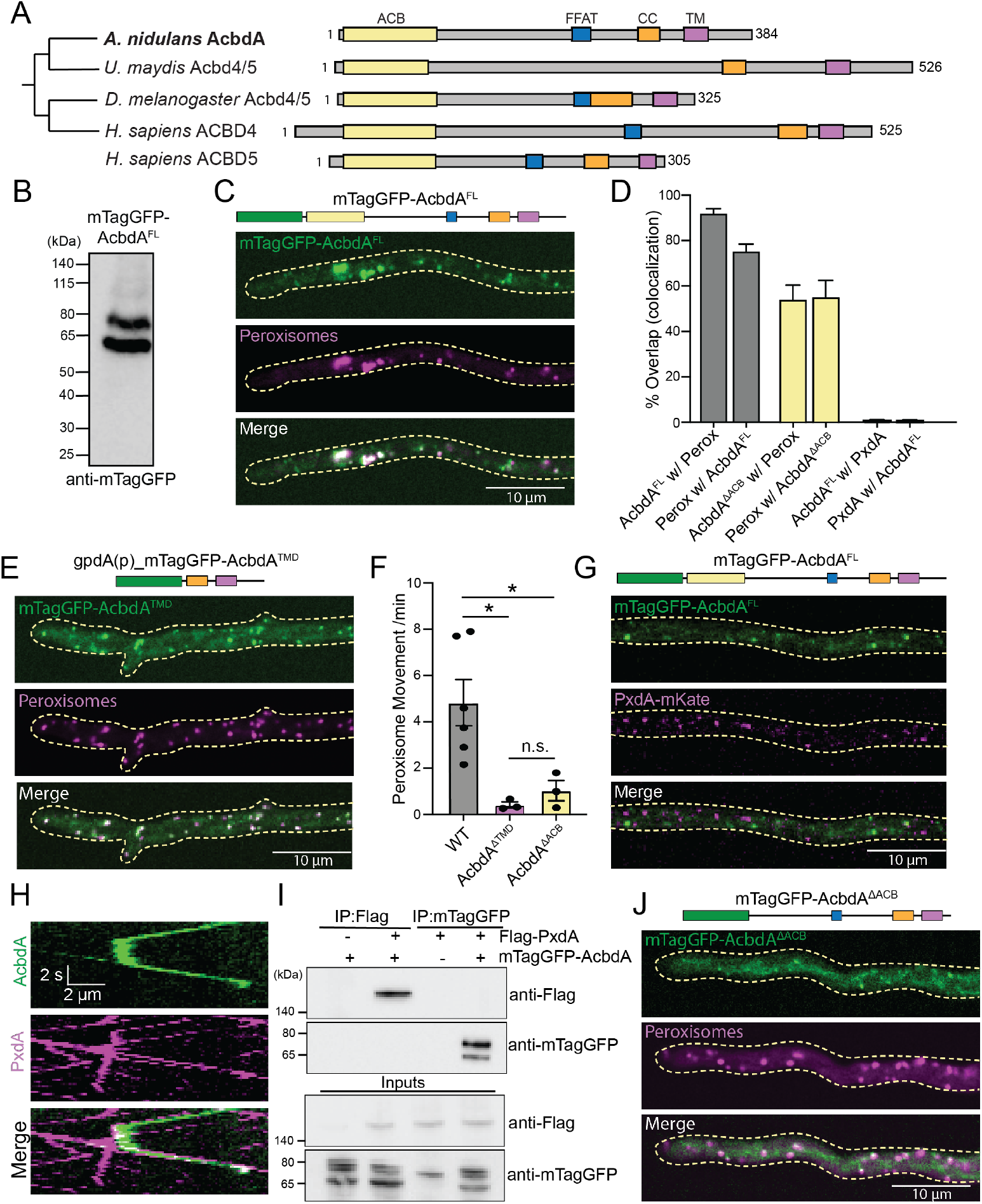
AcbdA localizes to peroxisomes via its transmembrane domain and requires its ACB domain for peroxisome hitchhiking on EEs. (**A**) Domain organization of *A.nidulans* AcbdA and other ACBD 4/5 homologs. ACB, FFAT, coiled-coil (CC) and transmembrane (TM) domains are highlighted. (**B**) Western blot N-terminally tagged mTagGFP-AcbdA^FL^. Predicted protein size is 71 kDa. (**C**) Domain organization of mTagGFP-AcbdA^FL^ construct (top) and representative micrographs of hyphae expressing mTagGFP-AcbdA^FL^ and mCherry-PTS1. (**D**) Bar graphs quantifying colocalization between mTagGFP-AcbdA^FL^/mCherry-Perox), mTagGFP-AcbdA^ΔACB^/mCherry-Perox, and mTagGFP-AcbdA^FL^/PxdA-mKate. Puncta were generated from the first frame of 20s two-color simultaneous image (n=27 (mTagGFP-AcbdA^FL^/mCherry-PTS1), n=33 (mTagGFP-AcbdA^ΔACB^/mCherry-PTS1), and n=21 (mTagGFP-AcbdA^FL^/PxdA-mKate) hyphae). (**E**) Domain organization of gpdA(p)_mTagGFP-AcbdA^TMD^ construct (top) and representative micrographs of hyphae expressing gpdA(p)_mTagGFP-AcbdA^TMD^ and mCherry-PTS1. (**F**) Bar graph of peroxisome movements per minute near the hyphal tip in hyphae expressing untagged constructs: AcbdA^FL^ (WT; see Methods), AcbdA^ΔACB^, and AcbdA^ΔTMD^. Mean peroxisome movements are 4.84 ± 0.99 for WT, 0.42 ± 0.12 for AcbdA^TMD^, and 1.03 ± 0.43 for AcbdA^ΔACB^ (*p<0.05; one-way ANOVA, Tukey’s post hoc multiple comparisons test; N=6 technical replicates from 65 hyphae (WT); N=3 technical replicates from 29 (AcbdA^ΔTMD^) and 30 (for AcbdA^ΔACB^) hyphae). (**G**) Representative micrographs of hyphae expressing mTagGFP-AcbdA^FL^ and PxdA-mKate. (**H**) Representative kymographs generated from a two-color movie of cotransporting mTagGFP-AcbdA^FL^ and PxdA-mKate. (**I**) Co-immunoprecipitation assay in HEK293 cells expressing AN_ mTagGFP-AcbdA^FL^ and AN_ PxdA-mKate. (**J**) Domain organization of mTagGFP-AcbdA^ΔACB^ construct (top) and representative micrographs of hyphae expressing mTagGFP-AcbdA^ΔACB^ and mCherry-PTS1. All error bars SEM.

All Acbd4/5 proteins characterized thus far have been demonstrated to associate with peroxisomes (Costello *et al*., 2017a, 2017b; Hua *et al*., 2017; Islinger *et al*., 2020; Kors *et al*., 2024). Therefore, we suspected that AcbdA may similarly localize to peroxisomes. To determine AcbdA localization, we endogenously tagged AcbdA with mTagGFP on the N-terminus. A lysate from this strain was resolved on SDS-PAGE followed by Western blot. mTagGFP-AcbdA resolves at a molecular weight consistent with its predicted molecular weight of ∼45 kDa plus the 27 kDa mTagGFP (Figure 3B).

mTagGFP-AcbdA was punctate along hyphae (Figure 3C). The vast majority of AcbdA puncta colocalize with peroxisomes (∼91%) and conversely, most peroxisomes colocalize with AcbdA (∼75%) (Figure 3D). Orthologs of AcbdA in the fungus *Ustilago maydis (*Um_Acbd4/5), in *Drosophila* (Dm_Acbd4/5), and in *H. sapiens* (Hs_ACBD4 and Hs_ACBD5) localize to peroxisomes via their TMDs (Costello *et al*., 2017a, 2023; Hua *et al*., 2017; Islinger *et al*., 2020; Kors *et al*., 2024). Similarly, AcbdA’s TMD is predicted to be a tail-anchored transmembrane domain targeted to peroxisomal membranes. To determine if AcbdA localizes to peroxisomes via its TMD, we developed a strain with GFP fused to the TMD (plus flanking regions) expressed from the exogenous gpdA promoter (gpdA-mTagGFP-AcbdA^TMD^). Almost all mTagGFP-AcbdA^TMD^ puncta colocalized with peroxisomes (Figure 3E), suggesting that the TMD is sufficient to drive peroxisome localization. Consistent with these findings, the mutagenized allele from our screen was a nonsense mutation located at the beginning of the TMD at aa W325 (Figure 1B) and deleting the TMD (AcbdA^ΔTMD^) abolished peroxisome movement (Figure 3F).

### AcbdA hitchhikes with PxdA on vesicles, but they do not interact in a heterologous system

We asked if the small subset of AcbdA puncta (∼9%) not colocalizing with peroxisomes were instead colocalized with PxdA-labeled EEs. We found no evidence of this (Figure 3, D and G). Therefore, it is likely that the very small pool of AcbdA either localizes to another organelle or we were unable to detect all instances of colocalization between AcbdA and peroxisomes due to our imaging conditions. It is also possible the remaining pool associates with Woronin bodies, a peroxisome derivative that has been previously shown to hitchhike on EEs (Songster *et al*., 2023).

Although no static AcbdA colocalized with PxdA, there were many instances of AcbdA puncta cotransporting and hitchhiking on motile PxdA-marked vesicles (Figure 3H, as an example). We asked if protein interactions between PxdA and AcbdA are responsible for hitchhiking at the peroxisome-EE MCS. To test this, we turned to a heterologous expression system by expressing Flag-PxdA alone, mTagGFP-AcbdA alone, or both in HEK-293 cells. Lysates from these cells were immunoprecipitated with either Flag- or GFP-conjugated beads. Western blot analysis indicated that although both proteins immunoprecipitated, neither PxdA nor AcbdA were present in coimmunoprecipitated lanes (Figure 3I). This finding suggests that PxdA and AcbdA do not directly interact. We cannot rule out that the PxdA-AcbdA interaction is too transient to identify with traditional immunoprecipitation methods. Furthermore, it is possible that these proteins are not capable of interacting in HEK-293 cells. Unfortunately, immunoblot signal from the mTagGFP-AcbdA strains (in *A. nidulans*) enriched by GFP-conjugated beads was barely detectable (Supplemental Figure S1D), precluding our ability to analyze AcbdA’s interaction with PxdA in *A. nidulans*.

### Acyl-coA binding domain is required for movement of peroxisomes

In addition to containing C-terminal TMDs, all Acbd4/5 proteins identified contain an N-terminal ACB domain (Islinger *et al*., 2020; Kors *et al*., 2024). Hs_ACBD5’s ACB domain is required for proper metabolism of VLCFAs (Costello *et al*., 2023). We were curious about the role of AcbdA’s ACB domain plays in peroxisome hitchhiking. We developed a strain with a deleted ACB domain (AcbdA^ΔACB^). Compared to WT, the AcbdA^ΔACB^ strain has drastically reduced peroxisome movement, phenocopying AcbdA^ΔTMD^-expressing strains (Figure 3F).

We predicted that AcbdA^ΔACB^ should still localize to peroxisomes since it has an intact TMD. Indeed, although colocalization and intensity of mTagGFP-AcbdA^ΔACB^ is reduced compared to mTagGFP-AcbdA^WT^, most puncta still localized to peroxisomes (Figure 3, D and J). We conclude that the ACB domain is not strictly required for peroxisome localization.

When analyzing peroxisome movement in our mTagGFP-AcbdA^FL^ strains, we noticed that they moved more frequently than WT (Supplemental Figure S1E). We hypothesized that this increase in peroxisome movement was a gain-of-function effect due to the presence of the GFP-tag near the ACB domain. Consistent with this, peroxisome movement was drastically reduced in mTagGFP-AcbdA^ΔACB^ compared to its full-length counterpart (Supplemental Figure S1E). Taken together, these data suggest that the ACB domain is critical for peroxisome hitchhiking.

### Fatty acid-induced movement of peroxisomes, but not proliferation, requires AcbdA

Given that AcbdA contains an ACB domain predicted to interact with acyl-coA activated FAs, we tested whether peroxisome hitchhiking is modulated by FAs. The movement of peroxisomes in WT strains was increased on acetate and butyrate compared to glucose (Figure 4A; Supplemental Figure S1F). We find that AcbdA is required for this FA-induced increase (Figure 4A; Supplemental Movie S3), suggesting that AcbdA and peroxisome hitchhiking is linked to FA metabolism.

**Figure 4.**
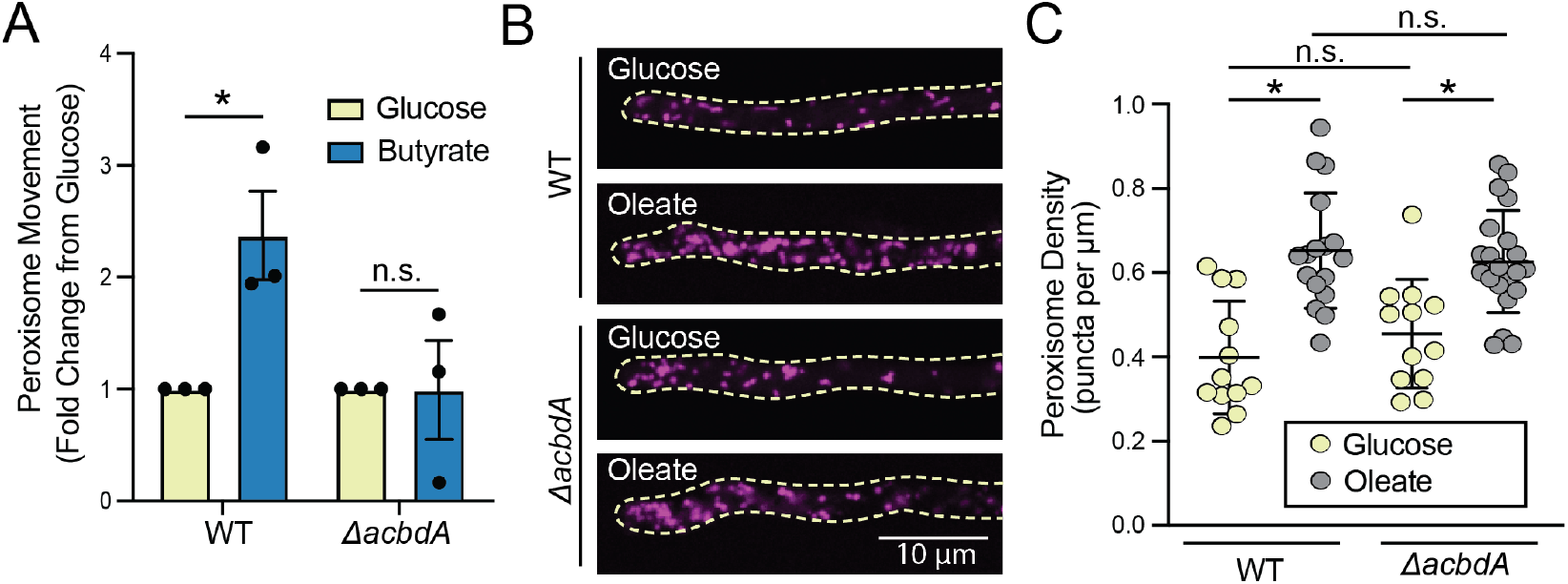
Fatty acid-induced movement of peroxisomes, but not number or proliferation, requires AcbdA. **(A)** Peroxisome movement quantified as a fold-change of 50 mM butyrate over 1% glucose conditions from each technical replicate (circles). Mean fold change is 2.37 ±0.39 (SEM) for WT and 0.99 ±0.44 (SEM) for *ΔacbdA* (*p<0.05; one-way ANOVA, Tukey’s post hoc multiple comparisons test; N=3 technical replicates from 30 (Glucose) and 33 (Butyrate) hyphae). Also see Supplemental Movie S3. (**B**) Representative micrographs of peroxisomes along WT and *ΔacbdA* hyphae grown on oleate (0.5% Tween-80) or 1% glucose. (**C**) Quantification of peroxisome density (puncta per hyphal distance in microns) for indicated conditions. Each circle represents individual hyphae. Mean ±S.D. are as follows: 0.40 ±0.13 (WT-Glucose), 0.65 ±0.14 (WT-Oleate), 0.46 ±0.13 *(ΔacbdA*-Glucose) and 0.63 ±0.12 *(ΔacbdA*-Oleate) (*p<0.01; one-way ANOVA, Tukey’s post hoc comparison; n.s.=not significant).

Previous studies demonstrated that FAs such as oleate induce peroxisome proliferation in *A. nidulans* (Valenciano *et al*., 1996, 1998; Hynes *et al*., 2008). To determine if AcbdA is required for FA-induced proliferation, we inoculated WT and *ΔacbdA* strains on either glucose or oleate as the sole carbon source and visualized peroxisomes (Figure 4B). A similar number of peroxisomes were present in WT and *ΔacbdA* hyphae grown on standard glucose media. When grown on oleate, peroxisome number was also increased, but to a similar extent in both WT and *ΔacbdA* hyphae, suggesting that AcbdA is dispensable for FA-induced proliferation (Figure 4C).

### The FFAT motif in AcbdA does not affect peroxisome movement

AcbdA does not regulate the baseline number or proliferation of peroxisomes. This is in contrast to other characterized Acbd4/5 proteins which have been shown to regulate the number of peroxisomes (Costello *et al*., 2017a; Kors *et al*., 2022, 2024). We were curious if other aspects of AcbdA were functionally distinct from its Acbd4/5 counterparts. Though all Acbd4/5 proteins interact with peroxisomes, Hs_ACBD4, Hs_ACBD5 and Dm_Acbd4/5 were previously shown to facilitate MCS between peroxisomes and the endoplasmic reticulum (ER) (Costello *et al*., 2017b, 2017a, 2023; Hua *et al*., 2017; Islinger *et al*., 2020; Kors *et al*., 2022, 2024). In both cases, ER localization is driven by a FFAT motif within the Acbd protein that binds to the integral ER membrane protein VAP (vesicle-associated membrane protein (VAMP)-associated protein) (Costello *et al*., 2017b, 2017a; Hua *et al*., 2017). Therefore, we sought to determine whether AcbdA contains a FFAT motif and whether the FFAT motif affects peroxisome motility. Using FFAT motif prediction tool (Murphy and Levine, 2016), we examined FFAT motif presence across the fungal kingdom. We found that, like the human and *Drosophila* Acbd4/5 homologs, AcbdA contains a FFAT motif (Figure 3A, Kors *et al*., 2024). Similar to previous studies, we identified FFAT motifs only in a subset of fungal species within the Pezizomycotina (Kors *et al*., 2024), and no FFAT motif within the fungi outside of the Pezizomycotina, including *U. maydis* Um_Acbd4/5, the only other characterized fungal Acbd protein (Figure 5A). This analysis suggests that ER association may be conserved in certain fungi, but not others (Kors *et al*., 2024).

**Figure 5.**
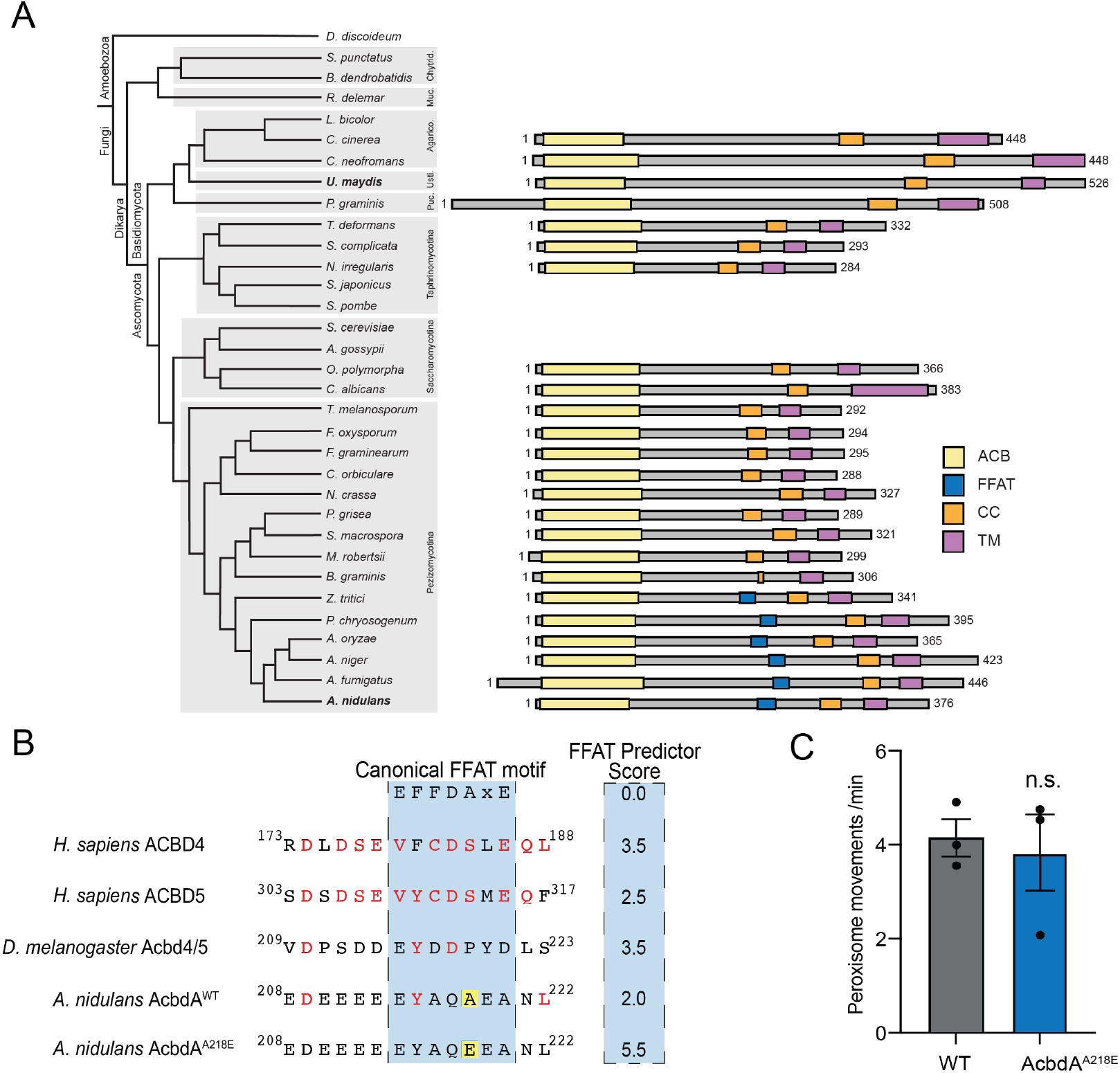
FFAT motif does not affect peroxisome movement. (**A**) Domain organization of ACBD protein homology among fungal species, including Pezizomycotina. (**B**) Sequence alignment against previously shown FFAT-motifs among different members of the Acbd4/5 family (Uniprot; FungiDB; Kors et al., 2024), shaded in blue. Conserved amino acids to Hs_ACBD4/5 are in red. Non-permissible A218E substitution is highlighted in yellow. FFAT predictor scores indicated. Scores ≤2.5 indicate a high probability of a functional FFAT while scores > 3.5 indicate improbability of a functional FFAT (Murphy and Levine, 2016). (**C**) Bar graph of peroxisome movements per minute near the hyphal tip in WT and AcbdA^A218E^ hyphae. Mean peroxisome movements are 4.16 ±0.40 for WT and 3.79 ±0.86 for AcbdA^A218E^ (p>0.9999 [n.s.]; Mann-Whitney test, N=3 technical replicates from 36 (WT) and 39 (AcbdA^A218E^) hyphae). All error bars SEM.

We then sought to determine whether the FFAT motif in AcbdA is critical for peroxisome movement. Mutating a critical residue at A218 to glutamic acid (A218E) had no affect on peroxisome movement, (Figure 5, B and C), suggesting that AcbdA’s FFAT motif is dispensable for peroxisome hitchhiking. We also fluorescently-labeled the ER (Sec63-GFP) (Markina-Iñarrairaegui *et al*., 2013); distribution was normal in *ΔacbdA* hyphae compared to WT (Supplemental Figure S1G). Together, these findings demonstrate that, though AcbdA has a FFAT motif like its human and *Drosophila* homologs, its FFAT motif is not critical for peroxisome movement.

### *A. nidulans* AcbdA has both similar and divergent properties to other Acbd4/5 proteins

Overall, we discovered AcbdA as a peroxisome-localized tether that interacts with EEs to facilitate peroxisome hitchhiking. We show that AcbdA localizes to peroxisomes via its TMD, and that its ACB domain, but not its FFAT motif, is critical for peroxisome hitchhiking. Finally, our data suggests that peroxisome hitchhiking is tied to FA-metabolism through AcbdA.

AcbdA displays functions that are in some cases consistent, and in others divergent, from known functions of other well-characterized Acbd4/5 proteins. Overall, AcbdA shares substantial similarity in domain organization with Hs_ACBD4, Hs_ACBD5, and Dm_Acbd4/5), as they all contain an ACB domain, FFAT motif, and TM domain (Figure 3A). Acbd4/5 proteins are known to facilitate MCS tethering. In humans, Acbd4/5 proteins assist in MCS formation between peroxisomes and the ER (Costello *et al*., 2017a, 2017b; Hua *et al*., 2017). On the other hand, we found that *A. nidulans* AcbdA facilitates MCS formation between peroxisomes and EEs. Furthermore, this MCS formation has differing effects on peroxisome motility. Hs_ACBD5 and Dm_Acbd4/5 are negative regulators of peroxisome movement and/or distribution (Costello *et al*., 2017a; Hua *et al*., 2017; Wang *et al*., 2018; Kors *et al*., 2024). Depleting Hs_ACBD5 increased peroxisome movement in human fibroblast cells and HeLa cells (Costello *et al*., 2017a; Hua *et al*., 2017). This is in contrast to AcbdA’s role in *A. nidulans*, where *ΔacbdA* strains displayed reduced peroxisome movement (Figure 2, C – E). Interestingly, overexpression of Hs_ACBD5 in hippocampal neurons reduced the movement and distribution of peroxisomes in the axon, but this was independent of FFAT interactions, suggesting that ER tethering may not play as critical a role in peroxisome mobility as previously thought (Wang *et al*., 2018). Future work will be needed to investigate the interplay between Acbd4/5 proteins’ FFAT interactions and peroxisome tethering to the ER under endogenous expression conditions, and how this affects peroxisome function.

One of the key similarities between AcbdA, and human Acbd4/5 are their ties to FA metabolism. Knockdown of Hs_ACBD5 reduces VLCFA metabolism and rescuing this defect requires its ACB domain (Costello *et al*., 2023). We found that the increase in peroxisome motility that occurs upon growth on FAs reqauires AcbdA in *A. nidulans*, suggesting a role for AcbdA in FA metabolism and its link to peroxisome motility. Overall, this is consistent with known functions of other Acbd4/5 proteins, which serve a dual function: MCS tethering and sequestering FAs, although future work will be needed to tease these mechanisms apart.

Interestingly, AcbdA functions appear to be more divergent from the other characterized fungal Acbd4/5 protein, *U. maydis* Um_Acbd4/5. Although *ΔUm_acbd4/5* strains had similar defects in peroxisome movement, they also displayed a drastic accumulation of EEs at the hyphal tip. This demonstrates that the peroxisome defects observed in *ΔUm_Acbd4/5* strains are likely not specific to peroxisome hitchhiking, and are a more general defect in vesicular trafficking. Curiously, Um_Acbd4/5 does not contain an FFAT motif and does not interact with VAP or facilitate contact with the ER (Kors *et al*., 2024).

Our data suggests that AcbdA does not directly interact with PxdA. Possible AcbdA interactors likely require the ACB domain, since removing it abolishes peroxisome movement. Future work will be required to determine the mechanisms by which AcbdA interacts with EEs to modulate peroxisome movement both in basal and FA-induced conditions.

## Abbreviations

ACB –: acyl-coA binding
DipA –: DenA interacting phosphatase
EE –: early endosome
ER –: endoplasmic reticulum
FA –: fatty acid
FFAT –: two phenylalanines in an acidic tract
MCS –: membrane contact site
PxdA –: peroxisome distribution mutant A
PTS1 –: peroxisome targeting signal 1
TMD –: transmembrane domain
VAP –: vesicle-associated membrane protein (VAMP)-associated protein
VLCFA –: very long-chain fatty acid
WT –: wild-type

## Acknowledgements

We would like to thank Michael J. Previs for helpful advice on figure illustrations. We would like to thank Samara Reck-Peterson (UCSD) and Reck-Peterson lab members Liv Songster, Gaurav Kumar, Patreece Suen, and Jessica Allen for helpful communication throughout the process. UT Health San Antonio Long School of Medicine and Zhao Lai for genomic library prep and whole genome sequencing. We thank Richard Todd (KSU) and Xin Xiang (USUHS) for helpful advice related to strain construction. Fungal Genetics Stock Center (KSU) for AN1062/AcbdA KO cassette and primers. Miguel Penalva (CSIC) for sec63-GFP (ER) strain. A.M.L. is supported by the Vermont Center for Immunobiology and Infectious Disease NIH T32 training grant 5T32AI055402-19. J.R.C. is supported by MOSAIC K99/R00 R00GM140269. J.S. is supported by R35GM150857 (NIH/NIGMS).

## Author contributions

B.D. and J.S. developed the project. B.D., M.B.F., R.A., A.M.L., M.B.O. and J.S. performed the wet lab experiments. I.G. and J.C. performed evolutionary analyses. B.D., J.R.C. and J.S. created illustrations.

B.D. J.R.C. and J.S. wrote the manuscript and all authors helped edit it.

## Movie Legends

**Movie S1. Peroxisome movement, but not EE movement is perturbed in AcbdA(mut) and PxdA (mut) found in mutagenesis screen**. Peroxisomes (magenta) and early endosomes (green) were imaged on 1% glucose media with inverted spinning-disk microscope (Nikon) using single-camera sequential acquisition.

**Movie S2. Peroxisome movement, but not PxdA-marked or DipA-marked EE movement is perturbed in *ΔacbdA* strains**. Peroxisomes (magenta), PxdA-marked EEs (green) and DipA-marked EEs were imaged on 1% glucose media with inverted spinning-disk microscope (Nikon) using single-camera sequential acquisition.

**Movie S3. Butyrate-induced increase in peroxisome movement is dependent on AcbdA**. Peroxisomes (magenta) were imaged on 1% glucose media and 50 mM butyrate media with inverted spinning-disk microscope (Nikon) using single-camera sequential acquisition.

## Materials and Methods

### Fungal Growth Conditions

*A. nidulans* strains were grown on yeast and glucose medium (YAG; [2% glucose, 0.5% yeast, trace elements (0.04 g/L sodium tetraborate, 0.4 g/L cupric sulfate, 1.0 g/L ferric orthophosphate, 0.6 g/L manganese sulfate, 0.8 g/L sodium molybdate, 8.0 g/L zinc sulfate)]; Szewczyk *et al*., 2006; Todd *et al*., 2007) or 1% minimal glucose medium ([1% glucose; trace elements, 0.052% MgSO_4_, Stock Salt (6 g/L NaNO_3_, 0.52 g/L KCl, 0.152 g/L KH_2_PO_4_)]; Nayak *et al*., 2006), both with 1% agar (BactoAgar). Media was supplemented with 1 mg/mL uracil, 2.4 mg/mL uridine, 2.5 µg/mL riboflavin, 1 µg/mL para-aminobenzoic acid, and 0.5 µg/mL pyridoxine when required (Nayak *et al*., 2006).

*A. nidulans* strains were also grown on minimal medium with 1% agar and with various fatty acids (50 mM sodium butyrate; 50 mM potassium acetate; 0.5% Tween-80 (oleate); Hynes *et al*., 2008) as the source of carbon substituted for the use of 1% glucose for applicable experiments. Due to slower growth on these fatty acid media, colonies were allowed to grow ∼24 hours.

For imaging of mature hyphae, a small number of spores were removed from a YAG growth plate with a pipette tip and inoculated onto minimal media agar plates supplemented with appropriate auxotrophs. Hyphae were allowed to mature for 18-22 hours at 37°C. Colonies on agar were excised and inverted onto Nunc Lab-Tek II 8-Chamber Coverglass plates for imaging. For imaging of germling hyphae, a saturating number of spores was inoculated in 0.01% Tween-80 and 0.2 µL/mL was added to liquid minimal media with appropriate supplements. A few different volumes (ranging from 250 to 500 µL) of liquid minimal media with spores were plated in Lab-Tek dishes and grown at 30°C for 18-22 hours. Chambers of equivalent confluence were selected for imaging. For all biochemistry (genomic DNA isolation and western blot), spores were inoculated in liquid yeast and glucose medium with supplements and allowed to grow 16-20 hours at 37°C either shaking at 220 rpm or in 100 mm petri dish.

### Plasmid and Strain Construction

Strains used and developed in this study are listed in Supplemental Figure S1A. All strains were confirmed by PCR/sequencing from genomic DNA (Lee and Taylor, 1990) and a combination of live-cell imaging and/or western blot analysis. Linear sequencing was performed by Plasmidsaurus using Oxford Nanopore Technology with custom analysis and annotation. For ΔacbdA strain construction, vector was obtained from Fungal Genetics Stock Center. For all other plasmids, inserts were annealed to the Blue Heron Biotechnology pUC vector at using Gibson isothermal assembly (Gibson *et al*., 2009). All plasmids were confirmed using Plasmidsaurus nanopore sequencing. Strains were created by homologous recombination at the endogenous locus with linearized DNA in strains lacking *ku70* with *Afribo (Aspergillus fumigatus ribo), AfpyrG (Aspergillus fumigatus pyrG), or Afpyro (Aspergillus fumigatus pyro)* as selectable markers (Straubinger *et al*., 1992; Nayak *et al*., 2006). New strains were created by introducing linearized DNA to protoplasts as described (Szewczyk *et al*., 2006), notably using an alternative lytic enzyme (Novozymes, VinoTastePro with 2500 PGNU/g Polygalacturonase and 75 BGXU/g Beta-glucanase). Additional strains were created using genetic crossing (Todd *et al*., 2007). All strains containing fluorescently labelled EEs and peroxisomes have been previously outlined (Egan *et al*., 2012; Tan *et al*., 2014; Salogiannis *et al*., 2016; Salogiannis, Christensen *et al*., 2021).

#### ΔacbdA strain

ANID_01062.1 genomic locus cassette and primers were procured from the Fungal Genetics Stock Center (McCluskey, 2003). Using forward primer: GTAACGCCAGGGTTTTCCCAGTCACGACGGTGGATCAGCTTCAGGATAC and reverse primer: GCGGATAACAATTTCACACAGGAAACAGCCCTCTACCCAGATTCAGAAG, we developed a knockout cassette containing an internal PyrG with 5’ 704 bp and 3’ 906 bp overhangs targeting the AN1062/AcbdA locus. *ΔacbdA* strain was validated by PCR using primers flanking the targeted region (Supplemental Figure 1B).

#### mTagGFP-AcbdA^FL^ strain

To visualize endogenously labelled AcbdA, a plasmid containing N-terminal mTagGFP-AcbdA was constructed. Codon-optimized mTagGFP succeeded by a GA(x4) linker was followed by endogenous *AN1062/acbdA* (including all introns), its native 3’UTR and an *Afribo* cassette and flanked by 1kb homologous recombination overhangs. To amplify the linearized DNA used for transforming protoplasts, the oligos 5’-CAATGCTTTCACGGGCGTCACCACG-3’ and 5’ CCGCTCCGTCCGGTAACTGAGGTCA-3’ were used. To allow for better targeting accuracy, the linear GFP-AcbdA-AfRibo-containing DNA into a ΔAcbdA strain (JSA 239; Supplemental Figure S1A) where the endogenous AcbdA locus was replaced with AfpyrG. Using this strategy, clones of interest were chosen that grew in the absence of appropriate selectable markers. This strain was confirmed by genotyping, PCR, imaging and western blot.

#### mTagGFP-AcbdA^ΔACB^strains

To visualize endogenously labelled AcbdA lacking the acyl-coA binding domain, a plasmid containing N-terminal GFP-AcbdA without amino acids 6-66 was constructed. Codon-optimized mTagGFP2 succeeded by a GA(x4) linker was followed by endogenous *AcbdA*^*ΔACB(6-66)*^ (incl. only intron 4), its native 3’UTR and an *Afribo* cassette and flanked by 1kb homologous recombination overhangs. To construct this, the insert 5’-GATCATATGGTTCTCCTGGAAAGTTTTTCGGCATGCTGCCACACTCATGGAATGGATGAACTTTACCGATCAGGTTTGCGTTCCCGTGCACAAGCATCAAATTCAGCGGTCGACGGTACAGCAGGTCCCGGTTCCACAGGCTCTCGCGGTGCAGGTGCAGGTGCAGGTGCAATGTCGGACTCTGTGGACGCCTGGTACGCTCAGCGTGGTTTATCCCGCACTGAGGCCAAACGGCGGTATATCACGACTCTCGTGGAGACAATGCACACCTACGCTTCGCAGACCGAGGAGGCGCGCGAGCTCGTCGCCGAACTTGAGTTTGTCTGGAATCAGGTGAAATCAAATATACCTTCGTCGACATCGAGCCCTGTGCAGTCCACGGGGGTTCCCCCGATTTCCCAACCGCAATCGCCGTATGGAAGTATAAGCGCGCAATTAGCACAGAACAATGAGTATCAGTATAAGACATCTACTGCGCGAGGAGACTCTCGGCTCCGTGTGTTGAGTCCCGTCAGTCAGCCAGATGATATTTATCAACGGCGTACGGCGCGGATGGGCTACGATCGTGATCAAGGGCTGGATCAAGGGGGTGACGATGAAAGTGTGAACTTAGACGAGGACGAAGAGGAGGAAGAATACGCCGAGGCCCAAGCCAATCTGTACGAGGACGATGATGAAGTGGAAGGTGAAGCAGGCGGCGCGGTCGATGAAGACGACGACGATGACCACCATCATCACCACCAACAGGTGTACTCGAGTCATATCCCGGATAATTCTCCAAGCCGGAAACGCGATCGTAAGCGCAACCACTACGGTAAAGACTCTTTTCCTCGCCAAACATGAGGAACTGACCTATTCTCATTCGAGAGACATGCTAATACAACGGGATACAGATGACATT GATAGCTGGCGATGGCGCCGCAGAGTCGAGCAGGCCTTGACCAAAATGACGGCCGAGATCGCCGCCGCCCGCGAGCAGATGGAAGCACGCACCCTGGCAGCCCGTCGAAGATCAGGCGTCTGGGCTTGGCTTCGGTGGCTGGTATGGGTCACCCTCCGACAGATTATCTGGGATCTGGCCCTCCTCGGCA TGTTATTGATCTGGATGCGGCTGCGCCAGGATCGACGGCTCGAGGAGCAGCTTAAGGCTGGGTGGTCGGTTGTGAAGGCGCGGTTAGCAGGACTGAAAGCTTTGCGGGACTTGAGGAAGGTATATATATTTTCTTGATTCTTATTCATCTTTTACTTTTACTCCCATCTGGTTTGGTTTCGTTTGTTCTTCACGAG TTCTATGCACGATAGTGCTTTCTATTCTGGTACATATTGACTAGATA-3’ was synthesized by Twist Bioscience and subcloned into *mTagGFP-AcbdA*^*FL*^ plasmid at 5’-SphI and 3’-AleI restriction enzyme cut sites. To amplify the linearized DNA used for transforming protoplasts, the oligos 5’-CAATGCTTTCACGGGCGTCACCACG-3’ and 5’ CCGCTCCGTCCGGTAACTGAGGTCA-3’ were used. This strain was confirmed by genotyping PCR followed by Plasmidsaurus nanopore sequencing as described above.

#### AcbdA-Ribo strain (Full-length control for untagged AcbdA domain analysis)

To construct this plasmid, site-directed mutagenesis was performed. Forward (5’-ATGTCGGACTCTGTG-3’) and reverse (5’-CATTGCGAATTAGAAAAGAC-3’) primers were developed using NEB Base Changer program. Linear DNA was generated from *mTagGFP-AcbdA*^*FL*^ by around the world PCR and re-annealed using T4 PNK and T4 DNA Ligase with the addition of ATP. To amplify the linearized DNA used for transforming protoplasts, the oligos 5’-CAATGCTTTCACGGGCGTCACCACG-3’ and 5’ CCGCTCCGTCCGGTAACTGAGGTCA-3’ were used. Strain was confirmed by linear DNA generated from genomic DNA isolation and sent to Plasmidsaurus.

#### AcbdA^ΔACB^ strain (untagged)

To construct this plasmid, site-directed mutagenesis was performed. Forward (5’-ATGTCGGACTCTGTG-3’) and reverse (5’-CATTGCGAATTAGAAAAGAC-3’) primers were developed using NEB Base Changer program to delete amino acids 6-66 (ACB). Linear DNA was generated from *mTagGFP-AcbdA*^*ΔACB*^ by PCR and re-annealed using T4 PNK and T4 DNA Ligase with the addition of ATP. To amplify the linearized DNA used for transforming protoplasts, the oligos 5’-CAATGCTTTCACGGGCGTCACCACG-3’ and 5’ CCGCTCCGTCCGGTAACTGAGGTCA-3’ were used. Strain was confirmed by linear DNA generated from genomic DNA isolation and sent to Plasmidsaurus.

#### AcbdA^ΔTMD^ (untagged)

To construct this plasmid, site-directed mutagenesis was performed. Forward (5’-ATGTCGGACTCTGTG-3’) and reverse (5’-CATTGCGAATTAGAAAAGAC-3’) primers were developed using NEB Base Changer program. Linear DNA was generated from *mTagGFP-AcbdA*^*ΔTMD*^ by PCR and re-annealed using T4 PNK and T4 DNA Ligase with the addition of ATP. To amplify the linearized DNA used for transforming protoplasts, the oligos 5’-CAATGCTTTCACGGGCGTCACCACG-3’ and 5’ CCGCTCCGTCCGGTAACTGAGGTCA-3’ were used. Strain was confirmed by linear DNA generated from genomic DNA isolation and sent to Plasmidsaurus.

#### gpdA(p)_mTagGFP-AcbdA^TMD^

To visualize the transmembrane domain of AcbdA, a plasmid containing N-terminal tagged AcbdA lacking the first 264 amino acids was generated from the gpdA promoter. The insert 5’-AAACGCGATCGTAAGCGCAACCACTACGATGACATTGATAGCTGGCGATGGCGCCGCAGAGTCGAGCAGGCCTTGACCAAAATGACGGCCGAGATCGCCGCCGCCCGCGAGCAGATGGAAGCACGCACCCTGGCAGCCCGTCGAAGATCAGGCGTCTGGGCTTGGCTTCGGTGGCTGGTATGGGTCACCCTCCGACAGATTATCTGGGATCTGGCCCTCCTCGGCATGTTATTGATCTGGATGCGGCTGCGCCAGGATCGACGGCTCGAGGAGCAGCTTAAGGCTGG GTGGTCGGTTGTGAAGGCGCGGTTAGCAGGAC TGAAAGCTTTGCGGGACTTGAGGAAGGTATATATATTTTCTTGA-3’ was synthesized and subcloned into a plasmid containing a gpdA promoter at the white locus by GenScript. To amplify the linear DNA used for transforming protoplasts, the oligos 5-CTCTGGAACAGTCTCGCCGTCTTG-3’ and 5’-ACTCGGAAAGGGTGGTGAATGCGG-3’ were used.

#### AcbdA^A218E^

FFAT sequence in *A. nidulans* was predicted from FFAT scoring algorithm (Murphy and Levine, 2016; Kors *et al*., 2024) to be EYAEAQA, preceded by a highly acidic tract: EDEEEE and the amino acids NL. The non-permissible substitution from A to E at the 5^th^ position of the FFAT motif (A218E) was determined from (Mikitova and Levine, 2012). Under this non-permissible substitution, the prediction score from the algorithm went from a 2.0 to a 5.5, functionally abolishing the FFAT motif (Murphy and Levine, 2016). To generate this plasmid, site-directed mutagenesis to target A218 was employed. Forward (5’-GGAAGAATACGCCGAGGAGCAAGCCAATCTGTACGA-3’) and reverse (5’-TCGTACAGATTGGCTTGCTCCTCGGCGTATTCTTCC-3’) were generated using NEB Base Changer program. DNA was amplified and annealed by PCR under high GC conditions from *AcbdA-Ribo* (full length control for untagged AcbdA domain analysis). To amplify the linearized DNA used for transforming protoplasts, the oligos 5’-CAATGCTTTCACGGGCGTCACCACG-3’ and 5’ CCGCTCCGTCCGGTAACTGAGGTCA-3’ were used.

### Fluorescence Microscopy and Image Acquisition

All microscopy was performed at 22°C using a Nikon Ti-2E inverted microscope with perfect focus, outfitted with a Yokogawa CSU-X1 spinning disk, two Photometrics Prime BSI sCMOS camera (TwinCam dual camera splitter with filter cubes), and four laser lines (405, 488, 561, and 647 nm). The microscope is controlled by a specialized computer that runs Nikon NIS-Elements. Equipped with three objectives: Plan Apo 20 × 0.75 NA, 60 × 1.4 NA, and 100 × 1.45 NA. All image acquisition was conducted with 60X oil objective. Peroxisome movies were generated with a 500 ms delay at a frame rate of 500 ms for 1 minute. Early endosome movies (RabA, PxdA, and DipA) were generated with no delay at a frame rate of 250 ms for 10 seconds. ER (sec63-GFP) was imaged at 100 ms exposure. For two-color near-simultaneous acquisition, mTagGFP-AcbdA^FL^ and either PxdA-mKate or mCherry-PTS1 were acquired at 30 ms exposures with no delay (60 ms frame rate) for 10 seconds.

### Data Analysis

All data were analyzed using FIJI/ImageJ (NIH). A blind analysis plug-in was used. To quantify movement of both EE-markers and peroxisomes, puncta were manually counted crossing a perpendicular line ∼10 µm from the hyphal tip. To generate kymographs, moving puncta were traced and resliced to create a kymograph using FIJI plug-in. Velocity and average run-length were calculated from the slope of generated kymographs. To easily quantify and identify legitimate runs, only events 4 frames (2 seconds) or greater were considered. To determine distribution from line-scan measurements, maximum-intensity projections of fluorescence and brightfield images were generated. Brightfield images were traced using a segmented line tool (width adjusted for width of hyphae), and traces were superimposed on fluorescence image. Average fluorescence intensity was projected over the area measured. Line-scans were normalized against baseline fluorescence intensity for each condition. Micrographs were deconvolved using an automatic deconvolution method with 50 µm pinhole size when indicated.

To correct for chromatic misalignment between channels for colocalization analysis, we used a Tetraspeck microsphere image as a reference. Channels 1 (green) and 2 (magenta) were isolated, and the Descriptor-based Registration (2D/3D) plugin in ImageJ/Fiji was used to align Channel 2 to Channel 1. Registration was performed using translational mode with the setting “approximately aligned” to guide initial feature matching. Alignment was based on intensity maxima of the fluorescent beads, with the detection threshold adjusted to ensure most puncta were identified. The resulting fused image was used to confirm alignment visually and saved for future reference. The alignment model derived from the Tetraspeck image was re-applied to experimental image stacks. This generated fused, registered images for each dataset, which were then used exclusively for subsequent colocalization analysis. LUTs were adjusted to standardize color representation across datasets (e.g., green for Channel 1, magenta for Channel 2), and brightness/contrast were normalized using the Auto function or consistent min/max values. For colocalization, puncta were manually identified in the first frame only and overlap fraction was determined.

Peroxisome number was determined by first subtracting background fluorescence with a Gaussian Blur. A mask of the ROI was created and maxima were then computed by FIJI. Maxima number was normalized to the length of the hyphae.

### 4NQO mutagenesis, genomic isolation, whole genome sequencing and variant analysis

Strains labelled with mCherry-FLAG-PTS1 and GFP-RabA (JSA69 strain; see Supplemental Figure S1A) were subjected to 4 Nitroquinoline 1-oxide (4NQO) mutagenesis described previously (Tan *et al*., 2014). Briefly, *A. nidulans* spores were diluted to 1×10^7^ in 0.1M KPO_4_ buffer with 0.01% Tween-80. Spores were subjected to 4NQO for 30 minutes at 37°C shaking at 200 rpm before quenching with 10% sodium thiosulfate. Spores were diluted 1:800 and plated onto YAG agar. Colonies were allowed to grow for 48-72 hours and survival rate was measured. For kill curve, spores were subjected to varying concentrations of 4NQO (see Supplemental Figure S1A). For mutagenesis screen, a final 4NQO concentration of 1.3 µg/ml was used.

For screen imaging, spores were inoculated and replica plated onto 1% agar minimal glucose media supplemented with appropriate auxotrophs. Colonies were allowed to grow for 16-20 hours at 37°C before colony edges were excised and inverted onto 96-well glass bottom plates (Cellvis) for imaging. All mutagenesis screening was conducted with the 20X objective.

Mutagenized strains of interest were backcrossed and progeny were pooled to create a library of genomic DNA. Mutagenized strains were crossed with a parent non-mutagenized strain and starved of oxygen and auxotrophs to promote sexual reproduction (Todd *et al*., 2007). Products of sexual reproduction, cleistothecia, were picked and cleaned after ∼7 days before rupturing in sterile water and plating diluted spores (to ensure separate, single colonies) onto YAG agar. Progeny were allowed to grow 48-72 hours before inoculating individual colonies onto minimal media plates with supplements. Progeny were screened and binned for presence of mutation and Mendelian genetic ratios were assured. Pools of progeny with and without mutated phenotype were created and used to co-inoculate liquid YAG media and allowed to grow 16-24 hours at 37°C in 100 mm petri dish. Mycelia was harvested, dried and ground with a mortar and pestle in liquid nitrogen for genomic DNA isolation.

For genomic DNA isolation, ground mycelia was subjected to lysis buffer (50 mM Tris-HCl, pH 7.2; 50 mM EDTA; 3% SDS; 1% 2-mercaptoethanol) at 65°C for 1 hour. Phenol::Chloroform was added and homogenous mixture was centrifuged (Fisherbrand accuSpin Micro17) at 10,000 x g for 15 minutes at 22°C. Aqueous phase was isolated before adding 3M sodium acetate and 0.54 volumes isopropanol and incubated for ∼30 minutes at -20°C. DNA solution was centrifuged at maximum speed for 3 minutes and washed with 70% ethanol. DNA pellet was resuspended in 50-100 µL of Tris-EDTA buffer (10 mM Tris, pH 8.0; 1 mM EDTA, pH 8.0).*

All sequence analysis was performed as previously described (Downes *et al*., 2014; Tan *et al*., 2014) using PartekFlow analysis software tools. After sequencing, a pre-alignment QA/QC score was determined. Data was aligned to the *A.nidulans* reference sequence (see www.aspergillusgenome.org/download/sequence/A_nidulans_FGSC_A4/current/A_nidulans_FGSC_A4_current_chromosomes.fasta.gz) using the Burrows-Wheeler Aligner (BWA). A post-alignment QA/QC score was determined. There was greater than 180-fold coverage of the 30 MB *A.nidulans* genome for all strains of interest and this included 99.67% representation of the entire genome. After Bam-filtering the data (>99.28% passed this filter), FreeBayes and GATK Haplotype variant analysis callers were used to determine variants compared to the reference genome. To identify putative causal alleles, synonymous mutations were removed, as well as nucleotide changes common to non-mutagenized parental strains and mutagenized, non-phenotypic strains. Only high-quality mutations (appearing greater than 5 times in mutagenized strains with the phenotype but absent from mutagenized non-phenotypic strains) were considered.

### Western blot and Immunoprecipitation Assay

All biochemistry was performed with ground mycelia as described above. For immunoprecipitation assay on *A. nidulans*, flash frozen samples were subjected to 3X mild lysis buffer (0.3% Triton X-100, 150 mM NaCl, 50 mM Tris pH 7.4, protease and phosphatase inhibitor) and allowed to nutate for 30 mins at 4°C. Lysates were then subjected to two consecutive max spins (Fisherbrand accuSpin Micro 17R) at 4°C, taking the supernatant from each. For immunoprecipitation assay on HEK-293 cells, cells were washed with PBS and lifted with cell lifters. Lifted cells were then treated to 1 mL of mild lysis buffer, followed by nutation at 4°C for 15 minutes and a max spin for 15 min at 4°C. Lysates for both species were treated to the same immunoprecipitation protocol. Lysates were rotated end-over-end in washed beads (Chromotek anti-GFP Trap Agarose, Sigma M2 anti-Flag affinity resin) for 1 hour at 4°C. Beads were washed 3 times with cold mild lysis buffer and sample buffer (2X; Laemmli, BioRad) was added. A western blot was performed. For western blot, flash frozen samples were subjected to 2X boiling urea buffer (8M Urea, 125 mM Tris-Cl, 1 mM EDTA, 2% SDS, 10 mM DTT, 4% glycerol) and incubated at 95°C for 15 minutes. Sample buffer was added. All protein samples were resolved on 4-12% gradient SDS-PAGE gels (NuPage) for 60-80 min at 150V. Gels were then transferred to nitrocellulose at 250 mA submerged in 1X transfer buffer for 3h at 4°C. Blots were blocked with 5% milk in TBS-0.1% Tween-20 (TBST-T). Antibodies were diluted in 3% BSA in TBS-T. Primary antibodies were incubated for ∼1 hour at RT and horseradish peroxidase (HRP)-conjugated secondary antibodies (Cell signaling technologies) were incubated for 30 min-1 hour at RT. Rabbit anti-Tag(CGY)FP (Evrogen) was used at 1:1000 to detect mTagGFP and mouse anti-DYDDDDK (Thermo Scientific) was used at 1:1000 to detect Flag. Blots were washed between all steps for 5-10 minutes with TBST-T. A ChemiDoc (BioRad) using Image Lab (3.0.1.14) software was used to image all blots.

### Domain Prediction

Homologs of ACBD4/5 proteins from organisms [*Aspergillus nidulans* FGSC A4 (NCBI:txid227321), *Aspergillus fumigatus* Af293 (NCBI:txid330879), *Aspergillus niger* CBS 513.88 (NCBI:txid425011), *Aspergillus oryzae* RIB40 (NCBI:txid510516), *Penicillium chrysogenum* (NCBI:txid5076), *Zymoseptoria tritici* ST99CH_3D1 (NCBI:txid1276537), *Blumeria graminis* (NCBI:txid34373), *Metarhizium anisopliae* ARSEF 23 (NCBI:txid655844), *Sordaria macrospora* (NCBI:txid5147), *Pyricularia grisea* (NCBI:txid148305), *Neurospora crassa* OR74A (NCBI:txid367110), *Colletotrichum orbiculare* MAFF 240422 (NCBI:txid1213857), *Fusarium graminearum* PH-1 (NCBI:txid229533), *Fusarium oxysporum* NRRL 32931 (NCBI:txid660029), *Tuber melanosporum* (NCBI:txid39416), *Candida albicans* (NCBI:txid5476), *Ogatea thermophila* (NCBI:txid460523), *Eremothecium gossypii* (NCBI:txid33169), *Saccharomyces cerevisiae* (NCBI:txid4932), *Schizosaccharomyces pombe* (NCBI:txid4896), *Schizosaccharomyces japonicus* (NCBI:txid4897), *Neolecta irregularis* (NCBI:txid48691), *Saitoella complicata* (NCBI:txid5606), *Taphrina deformans* (NCBI:txid5011), *Puccinia graminis* (taxid:5297), *Ustilago maydis* (NCBI:txid5270), *Cryptococcus neoformans* (NCBI:txid5207), *Coprinopsis cinerea* (taxid:5346), *Laccaria bicolor* (taxid:29883), *Rhizopus delemar* (NCBI:txid936053), *Batrachochytrium dendrobatidis* (NCBI:txid109871), *Spizellomyces punctatus* (NCBI:txid109760), *Dictyostelium discoideum* (NCBI:txid44689), *Homo sapiens* (NCBI:txid9606), *Drosophila melanogaster* (NCBI:txid7227)] were identified using a combination of the Basic Local Alignment Search Tool (BLAST) (Altschul *et al*., 1990) and literature review. We used standard BLASTp parameters for all searches (max target sequences = 5000, E = .05, word size = 5, BLOSUM 62 matrix). Multiple splice variants were not included.

ACBD4/5 homologs were initially identified using the *A. nidulans* full length protein sequence as a BLASTp query. Follow-up BLASTp searches used full length protein sequences from *N. irregularis, C. orbiculare, U. maydis*, and *A. gossypii* to identify additional homologs, and reciprocal searches were performed in all cases. To classify a protein as an ACBD4/5 homolog, it was required to have an ACBD domain, coiled-coil domain, and transmembrane (TM) domain. ACB domains were identified using SMART (Schultz *et al*., 2000) or NCBI protein entries. TM domains were predicted using DeepTMHMM (Hallgren *et al*., 2022), TMHMMv2 (Sonnhammer *et al*., 1998; Krogh *et al*., 2001), and AlphaFold Server prediction (Abramson *et al*., 2024). Coiled-coil domains were predicted using CoCoNat (Madeo *et al*., 2023) and WaggaWagga algorithm Ncoils (Simm *et al*., 2015). FFAT motifs were predicted using a published calculator (Murphy and Levine, 2016). Only FFAT motif scores below 2.5 were included in our diagrams. Protein schematics were created using drawProteins (Brennan, 2018) and finalized in Adobe Illustrator 2025.

**Figure S1.**
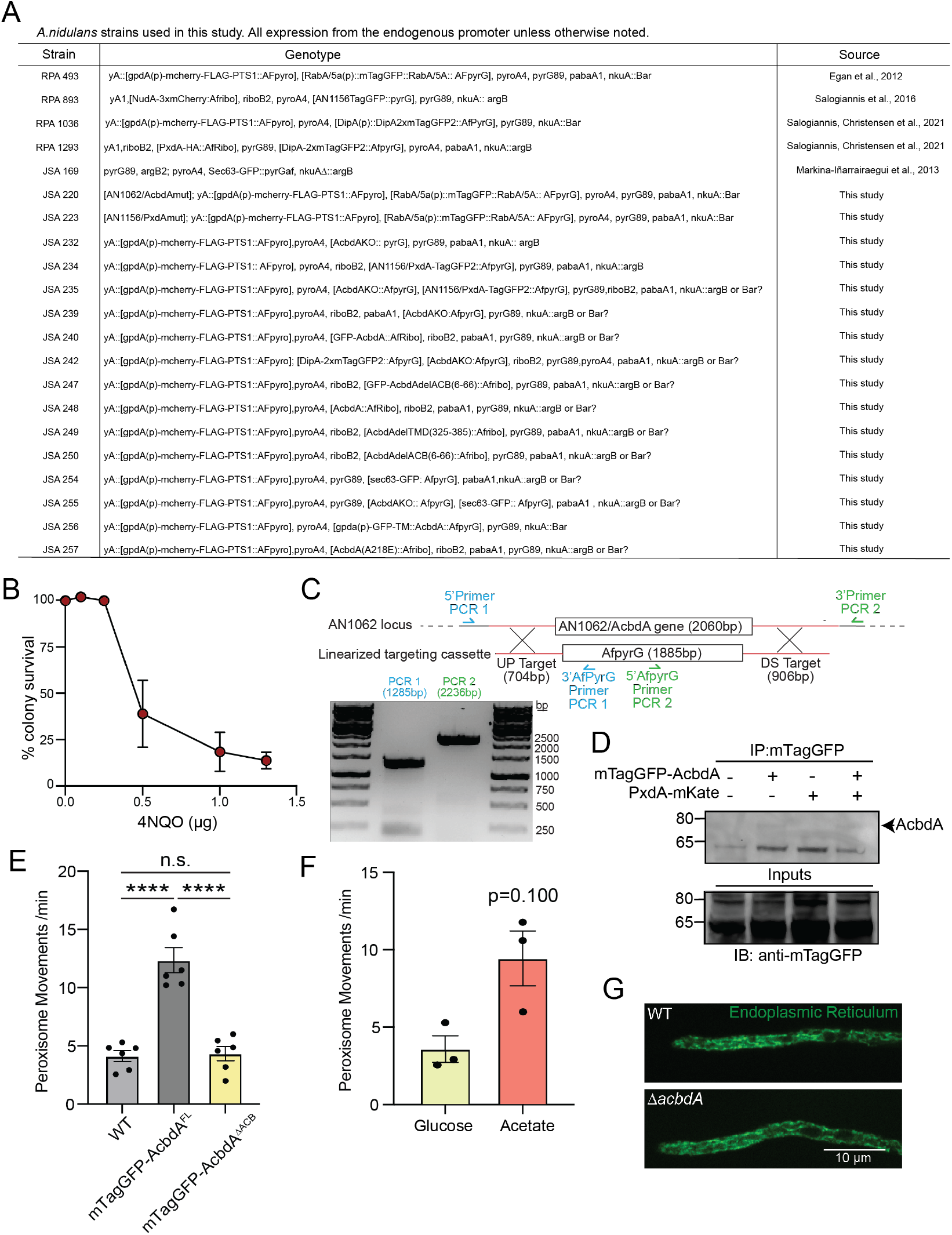
Characterization of AcbdA. (**A**) Table of all *A. nidulans* strains used in this study. (**B**) Survival rate curve of 4NQO mutagenesis under increasing concentrations (µg). (**C**) Verification of *ΔacbdA* by PCR. PCR fragment 1 (blue) is lane 1 and PCR fragment (green) is lane 2. (**D**) Immunoprecipitation (IP:mTagGFP) of strains expressing endogenous mTagGFP-AcbdA and PxdA-mKate. (**E**) Bar graph of peroxisome movements per minute in WT, mTagGFP-AcbdA^FL^, and mTagGFP-AcbdA^ΔACB^ strains. Mean peroxisome movements are 4.14 ±0.45 for WT, 12.36 ±1.07 for mTagGFP-AcbdA^FL^ and 4.35 ±0.61 for mTagGFP-AcbdA^ΔACB^ (****p<0.0001; one-way ANOVA, Tukey’s multiple comparisons test, N=6 technical replicates from 72 (WT), 60 (mTagGFP-AcbdA^FL^), and 63 (mTagGFP-AcbdA^ΔACB^) hyphae). (**F**) Bar graph of peroxisome movements per min in WT strains under 1% glucose and 50 mM acetate conditions. Mean peroxisome movements are 3.61±0.85 for 1% glucose and 9.46 ±1.76 for 50 mM acetate (p=0.100; Mann-Whitney test, N=3 technical replicates from 38 (1% glucose) and 29 (50 mM Acetate) hyphae). (**G**) Representative micrographs of Sec63-GFP (endoplasmic reticulum) in WT and *ΔacbdA* strains. All error bars SEM.

